# ENSO drives lateral separation of FAD-associated skipjack and bigeye tuna in the Western Tropical Pacific

**DOI:** 10.1101/2020.06.25.170613

**Authors:** Shirley Leung, K. A. S. Mislan, LuAnne Thompson

## Abstract

Incidental capture of juvenile bigeye tuna in fish aggregating device (FAD)-associated purse seine fisheries targeting skipjack tuna have contributed significantly to the degradation of bigeye stocks in the Western Tropical Pacific (WTP) Ocean. One way to reduce this incidental catch is to simply limit purse seine fishing effort; however, skipjack tuna stocks are healthy and economically important to many small island nations in the region. Here we assess whether there is sufficient lateral separation of skipjack and bigeye within FAD-associated purse seine fisheries in the WTP to allow limiting bigeye catch while maintaining a robust skipjack fishery. Based on monthly 5°-longitude-by-5°-latitude catch and effort data, FAD-associated bigeye and skipjack catch per unit effort (CPUE) covary tightly throughout the WTP, such that lateral separation between the two species is generally small. There are, however, significant variations in the amount of separation over both space and time. Waters within the Party to the Nauru Agreement exclusive economic zones (EEZs) belonging to Palau, Solomon Islands, and Tuvalu regularly exhibit some of the smallest bigeye-to-skipjack catch ratios, especially during El Niño. In contrast, waters within Kiribati’s Phoenix Islands EEZ regularly exhibit some of the largest bigeye-to-skipjack catch ratios, which are particularly high during La Niña. In general, El Niño lowers bigeye-to-skipjack catch ratios east of 170°E, while La Niña lowers bigeye-to-skipjack catch ratios west of 170°E. These ENSO-driven variations in separability are larger and more widespread than those driven by seasonality, due to larger associated variations in environmental conditions. Sea surface height anomalies may be particularly useful for demarcating the different environments preferred by skipjack and bigeye throughout the WTP. Sea surface temperatures, temperatures at 100 m, and thermocline depths may also help distinguish between the two species’ preferred habitats in many areas. These analyses can help better inform the complex decisions made by both fishers during operations and fisheries managers during creation of effective, dynamic policies to preserve bigeye stocks in the WTP. They also show that climate variability can have substantial effects on the spatial distributions of top pelagic predators and their interactions with one another.

## 1 Introduction

The Western and Central Pacific Ocean (WCPO) accounts for more than 60% of total global tuna catch [1]. Tuna and other highly migratory species in this region are managed by the Western and Central Pacific Fisheries Commission (WCPFC), which is one of five tuna Regional Fishing Management Organizations in operation around the world. The most commonly fished tuna species in the WCPO are skipjack (*Katsuwonus pelamis*), yellowfin (*Thunnus albacares*), bigeye (*Thunnus obesus*), and albacore tuna (*Thunnus alalunga*) [2]. Though these sought-after tuna species often inhabit the same waters within the WCPO, they historically have had very different stock statuses. For example, skipjack stocks have always been considered healthy [3,4], while bigeye stocks were considered overfished and undergoing overfishing until the most recent stock assessment in 2017 [5–7]. Differences between the 2017 stock assessment and those that came before were due to use of an updated growth model calibrated to new otolith age data [6– 9]. Based on the 2017 stock assessment, the WCPFC adopted a tropical tuna bridging measure that increased bigeye catch limits for WCPO longline fisheries by 11% [10]. A follow-up analysis, however, indicated that there is a >20% risk of bigeye populations falling below the limit reference point (adult biomass relative to unfished levels = SB/SB_F=0_ = 0.2) over the next 30 years with this bridging measure in place [11,12], leading some scientists and conservationists to oppose the measure [13,14]. In response to these concerns, the WCPFC encouraged more “research to identify ways for purse seine vessels to minimize the mortality of juvenile bigeye tuna” [10].

Incidental take of juvenile bigeye tuna in fish aggregating device (FAD)-associated purse seine fisheries has indeed contributed significantly to bigeye stock depletion in the WCPO [15– 17]. The simplest way to reduce the number of juvenile bigeye caught in purse seine sets is to limit effort, but skipjack tuna caught by purse seine are not at risk of being overfished and have tremendous economic value for small island nations in the region [1,3,4]. The research efforts encouraged by the WCPFC should thus focus on reducing incidental juvenile bigeye catch while maintaining current levels of purse seine skipjack catch. Past work within the Tropical Pacific has indeed taken this constraint into account [18–23]. For instance, Harley & Suter (2007) [18] investigated whether time-area closures in the Eastern Pacific Ocean could reduce purse seine bigeye catch without also significantly reducing that of skipjack. They found that in the best case scenario, a 3-month closure in the Eastern Equatorial Pacific during the third quarter of the year could reduce bigeye catch by 11.5%, while reducing skipjack catch by 4.3%. This led them to conclude that because of the strong tendency for bigeye and skipjack to be caught together, reducing bigeye catches further would not be possible via time-area closures without unacceptably large reductions in skipjack catch. Using ultrasonic telemetry, Schaefer & Fuller (2013) [23] documented the simultaneous behaviors of skipjack, bigeye, and yellowfin tuna within large, FAD-associated aggregations in the Eastern Equatorial Pacific; they found that differences in the behavior and vertical distributions of the three tuna species were not big enough that practical modifications to purse seine fishing practices could effectively avoid capturing small yellowfin and bigeye tunas while still optimizing skipjack capture. Leroy et al. (2009) [21] also concluded that the potential for targeting specific species through fishing depth selection is limited, based on an analysis of the vertical behaviors of the same three species associated with anchored FADs within Papua New Guinea’s exclusive economic zone (EEZ). Given the difficulties associated with separating bigeye and skipjack vertically and/or via time-area closures, Hu et al. (2018) [19] sought to separate the two species laterally by investigating differences in their habitat preferences using 1°-latitude by 1°-longitude monthly purse seine catch and effort data in the Eastern Tropical Pacific (ETP). They found that within the ETP, purse seine-caught bigeye occupy waters further from the coast where the hypoxic layer is deeper, while purse seine-caught skipjack occupy more productive waters associated with equatorial and coastal upwelling. Thus, to reduce incidental bigeye catch rates, Hu et al. (2018) [19] suggested targeting skipjack where hypoxic layers are shallowest (i.e., where there is coastal upwelling).

Here we expand upon this previous work by analyzing lateral FAD-associated, purse seine skipjack-bigeye catch separability in the Western Tropical Pacific (WTP), where the majority of skipjack tuna are caught [2]. Oceanographic conditions are quite different in the WTP compared to the ETP regions examined by Harley & Suter (2007) [18], Hu et al. (2018) [19], Lennert-Cody et al. (2008) [20], and Schaefer & Fuller (2013) [23] (e.g., [24,25]). Furthermore, new growth rate measurements suggest that Tropical Pacific bigeye potentially consist of two separate stocks (though with mixing in between), divided between the WTP and ETP at approximately 150°W [9,13]. Incidental catches of juvenile bigeye may therefore have quite different patterns and causes in the WTP compared to neighboring regions, necessitating WTP-specific analyses of lateral skipjack-bigeye separability. Previous studies have also overlooked the potentially important effects of El Niño Southern Oscillation (ENSO) on skipjack-bigeye purse seine catch separability. Rather, they have focused primarily on seasonal effects (i.e., [18]) or on ENSO-related variations in purse seine skipjack catch alone (e.g., [26,27]). Studies quantifying ENSO-driven variations in bigeye catch have focused solely on adult longline fisheries [28,29], and have not examined incidental juvenile purse seine capture.

Here we quantify the effects of both seasonality and ENSO on lateral skipjack-bigeye catch separability within FAD-associated purse seine fisheries in the Western Tropical Pacific. We also examine the environmental conditions that drive lateral separation of skipjack and bigeye catches in this region. Based on these analyses, we offer suggestions on how to maximize skipjack catch while simultaneously minimizing incidental juvenile bigeye catch within FAD-associated purse seine fisheries in the WTP.

## 2 Materials and methods

### 2.1 Data sources

#### 2.1.1 Purse seine catch and set data

Public domain purse seine fisheries catch and effort data was obtained at https://www.wcpfc.int/folder/public-domain-data on May 1, 2019 (date of issue: July 18, 2018). This data was aggregated by the WCPFC from information gathered in fishing vessel logbooks and by observers. The temporal resolution is monthly and the spatial resolution is 5°-latitude by 5°-longitude. Any monthly 5°-by-5° grid cell with data from less than three vessels was excluded to protect operational privacy [30]. This dataset runs from January 1967 to December 2017, though not all variables are available throughout this entire period. Catch (in metric tons) is broken up by species (skipjack, yellowfin, bigeye, and other), while sets are broken up by type (unassociated schools, natural log/debris, drifting FAD, anchored FAD, and other). S1 Fig shows the total number of sets of each type. In this study, we examine only FAD-associated skipjack and bigeye catches (i.e., from sets made on natural log/debris, drifting FADs, and anchored FADs only). For a given month and grid cell, we compute skipjack and bigeye catch per unit effort (SKJ and BET CPUE, respectively) by dividing skipjack and bigeye catch by the total number of FAD-associated sets. We assume that CPUEs are a reasonable proxy for abundance. We also compute bigeye-to-skipjack (BET:SKJ) catch ratios by dividing SKJ CPUE by BET CPUE.

#### 2.1.2 Oceanographic data from in situ observations

*In situ* profiles of temperature, salinity, and dissolved oxygen concentrations (O_2_) were downloaded from the World Ocean Database (WOD) at https://www.nodc.noaa.gov/OC5/SELECT/dbsearch/dbsearch.html on December 9, 2018. WOD gathers and stores quality-controlled data from buoys, ships, gliders, and floats [31]. Flagged profiles were excluded and XBT/MBT temperatures were corrected using Levitus et al. (2009) [32]. The profiles of temperature, salinity, and O_2_ down to 700 m depth were binned onto the same 5°-by-5° grid as the WCPFC data and then averaged over each month within the WCPFC data’s timeframe (Jan 1967 - Dec 2017). These monthly mean maps were then used to calculate corresponding maps of thermocline depth, oxygen partial pressure (pO_2_), and tuna hypoxic depth. Thermocline depths were computed using the variable representative isotherm method, as recommended for tropical waters by Fiedler (2010) [33]. To describe oxygen availability, we report water column oxygen content in terms of both partial pressures (pO_2_) as well as dissolved concentrations (O_2_). Though dissolved concentrations are more commonly used, pO_2_ is more biologically relevant as it provides the force which drives oxygen transfer into animal tissue [34,35]. To compute pO_2_ from input variables of temperature, salinity, and O_2_, we convert O_2_ to percent oxygen saturation [36], divide percent oxygen saturation by the fractional atmospheric concentration of oxygen (21%), and then correct for hydrostatic pressure at depth [37]. To describe oxygenated vertical habitat availability, we use tuna hypoxic depth, defined as the shallowest depth at which pO_2_ first falls below 15 kPa [25]. This 15 kPa pO_2_ threshold is approximately equivalent to the 3.5 ml l^-1^ dissolved concentration threshold below which conditions are considered hypoxic for skipjack tuna [38–44].

#### 2.1.3 Oceanographic data from satellite and reanalysis

Monthly mean mixed layer depths interpolated to a 0.5°-by-0.5° grid were obtained from ECCO Version 4, Release 4 [45,46] global ocean state estimate (also known as ECCO reanalysis) at https://ecco.jpl.nasa.gov/drive/files/Version4/Release4/interp_monthly/MXLDEPTH on March 12, 2020. Data coverage is from January 1992 – December 2017.

Monthly mean chlorophyll-a concentrations interpolated to a 4-by-4 km grid were obtained from the European Space Agency Ocean Colour Climate Change Initiative version 4.2 [47] at ftp://oc-cci-data:ELaiWai8ae@ftp.rsg.pml.ac.uk/occci-v4.2/geographic/netcdf/monthly/chlor_a/ on March 12, 2020. Data coverage is from September 1997 to December 2017.

Monthly mean sea surface height anomalies interpolated to a 0.25°-by-0.25° grid were obtained from AVISO+ at https://www.aviso.altimetry.fr/en/data/products/sea-surface-height-products/global/gridded-sea-level-anomalies-mean-and-climatology.html on October 18, 2019. Data coverage is from January 1993 to December 2017.

After downloading, all monthly mean satellite maps were re-interpolated onto the same 5°-by-5° grid as the WCPFC data. S2 Fig shows the total number of months of data available for each variable discussed in Section 2.1, while Table 1 summarizes pertinent information about each variable.

**Table 1.**
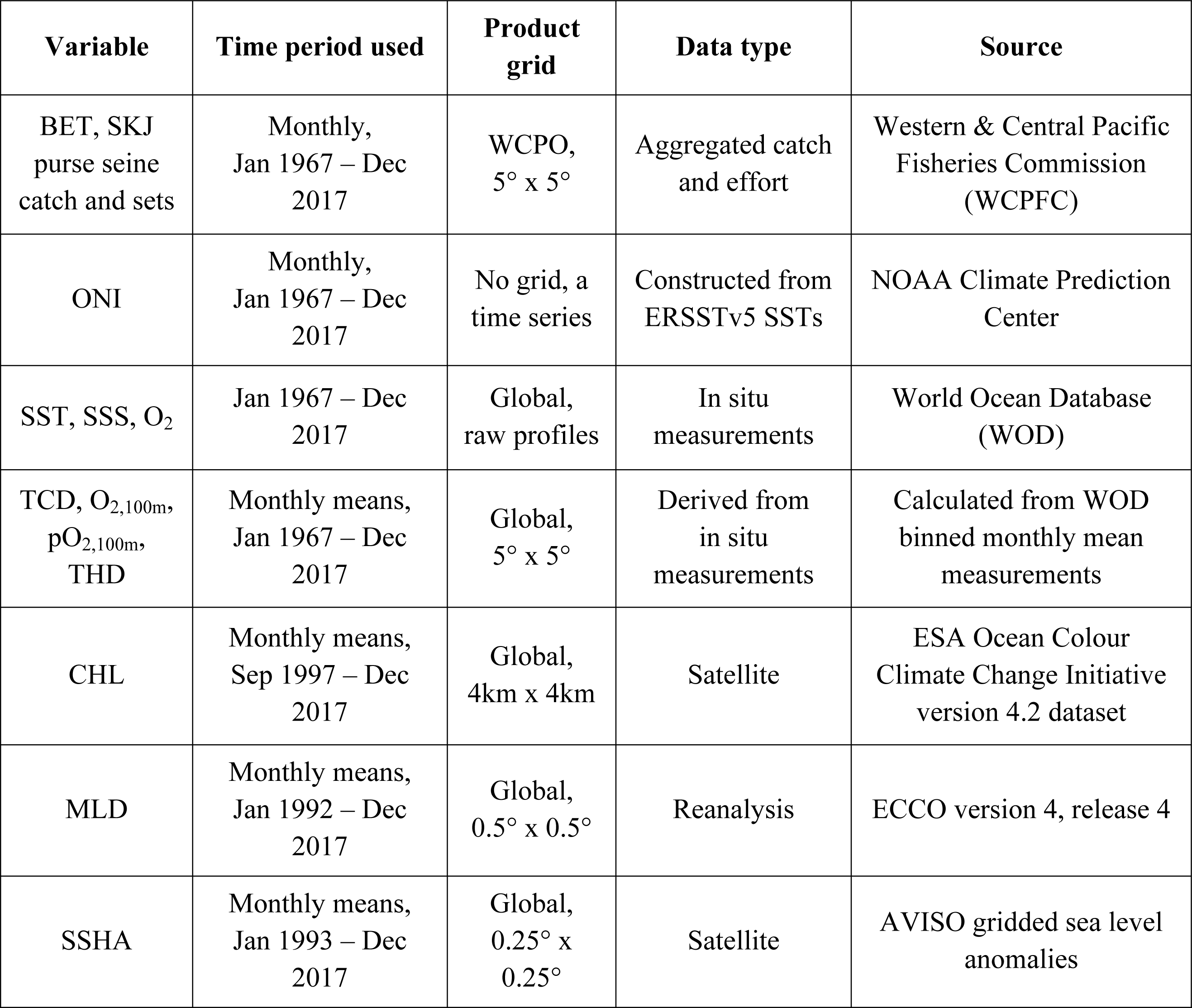
Summary of datasets utilized.

| Variable | Time period used | Product grid | Data type | Source |
| --- | --- | --- | --- | --- |
| BET, SKJ<br>purse seine<br>catch and sets | Monthly,<br>Jan 1967 – Dec<br>2017 | WCPO,<br>5° x 5° | Aggregated catch<br>and effort | Western & Central Pacific<br>Fisheries Commission<br>(WCPFC) |
| ONI | Monthly,<br>Jan 1967 – Dec<br>2017 | No grid, a<br>time series | Constructed from<br>ERSSTv5 SSTs | NOAA Climate Prediction<br>Center |
| SST, SSS, O<sub>2</sub> | Jan 1967 – Dec<br>2017 | Global,<br>raw profiles | In situ<br>measurements | World Ocean Database<br>(WOD) |
| TCD, O<sub>2,100m</sub>,<br>pO<sub>2,100m</sub>,<br>THD | Monthly means,<br>Jan 1967 – Dec<br>2017 | Global,<br>5° x 5° | Derived from<br>in situ<br>measurements | Calculated from WOD<br>binned monthly mean<br>measurements |
| CHL | Monthly means,<br>Sep 1997 – Dec<br>2017 | Global,<br>4km x 4km | Satellite | ESA Ocean Colour<br>Climate Change Initiative<br>version 4.2 dataset |
| MLD | Monthly means,<br>Jan 1992 – Dec<br>2017 | Global,<br>0.5° x 0.5° | Reanalysis | ECCO version 4, release 4 |
| SSHA | Monthly means,<br>Jan 1993 – Dec<br>2017 | Global,<br>0.25° x<br>0.25° | Satellite | AVISO gridded sea level<br>anomalies |

#### 2.1.4 ENSO index

The ENSO index used here, called the Oceanic Niño Index or ONI, is calculated as the three-month running mean of sea surface temperature anomalies in the Niño 3.4 region (5°N-5°S, 120°-170°W) [48]. Periods when ONI is greater than 0.5°C or less than −0.5°C for at least five consecutive three-month running-mean periods are classified as El Niño and La Niña phases, respectively. Time series of ONI were downloaded from http://origin.cpc.ncep.noaa.gov/products/analysis_monitoring/ensostuff/ONI_v5.php on June 4th, 2018.

### 2.2 Spatiotemporal computations

#### 2.2.1 Fieldwise statistical significance

To avoid incorrect and/or overstated interpretations, we control the fieldwise false discovery rate (FDR) for each set of multiple hypothesis tests conducted here (Figs 1, 4c-d, 5-6, 8, c-d in S3-4 Figs, column 2 in S5-7 Figs, and S10-12 Figs). The FDR is the statistically expected fraction of local null hypothesis rejections for which the null hypotheses are actually true. In this context, local is defined as being at a single grid point in the case of mapped analyses (Figs 1, 4c-d, 8, c-d in S3-4 Figs, column 2 in S5-7 Figs, S10-12 Figs) or within a single EEZ in the case of comparisons across EEZs (Figs 5-6). Controlling the FDR in the context of multiple-hypothesis tests requires smaller *p*-values to reject local null hypotheses compared to a single-hypothesis test. The procedure for determining significance under a specified FDR is as follows [49]: 1.) Sort the collection of *p*-values from N (i.e., the total number of grid cells or total number of EEZs) local hypothesis tests *p*_*i*_, with *i* = 1, …, N, in ascending order. 2.) Reject local null hypotheses only if their respective *p*-values are smaller than or equal to a threshold level *p*_*FDR*_, where *p*_*FDR*_ is equal to the largest *p*_*i*_ that is smaller than or equal to the fraction of *α*_*FDR*_ specified by *i/N. p*_*FDR*_ is thus calculated as follows:

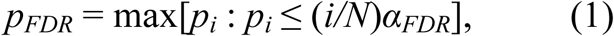

where *α*_*FDR*_ is the chosen FDR (expressed as a fraction). This method assumes that the multiple local tests are statistically independent, but is also valid when the results of the multiple tests are strongly correlated, as is the case with the spatially and temporally autocorrelated fisheries and oceanographic data used here. Indeed, for data of this nature, the achieved FDR will be smaller than the specified FDR by about two times; the specified FDR should thus be approximately double the desired level [49–51]. In this study, we specify an FDR of 0.1 for all analyses, resulting in an achieved FDR of ∼0.05.

**Fig 1.**
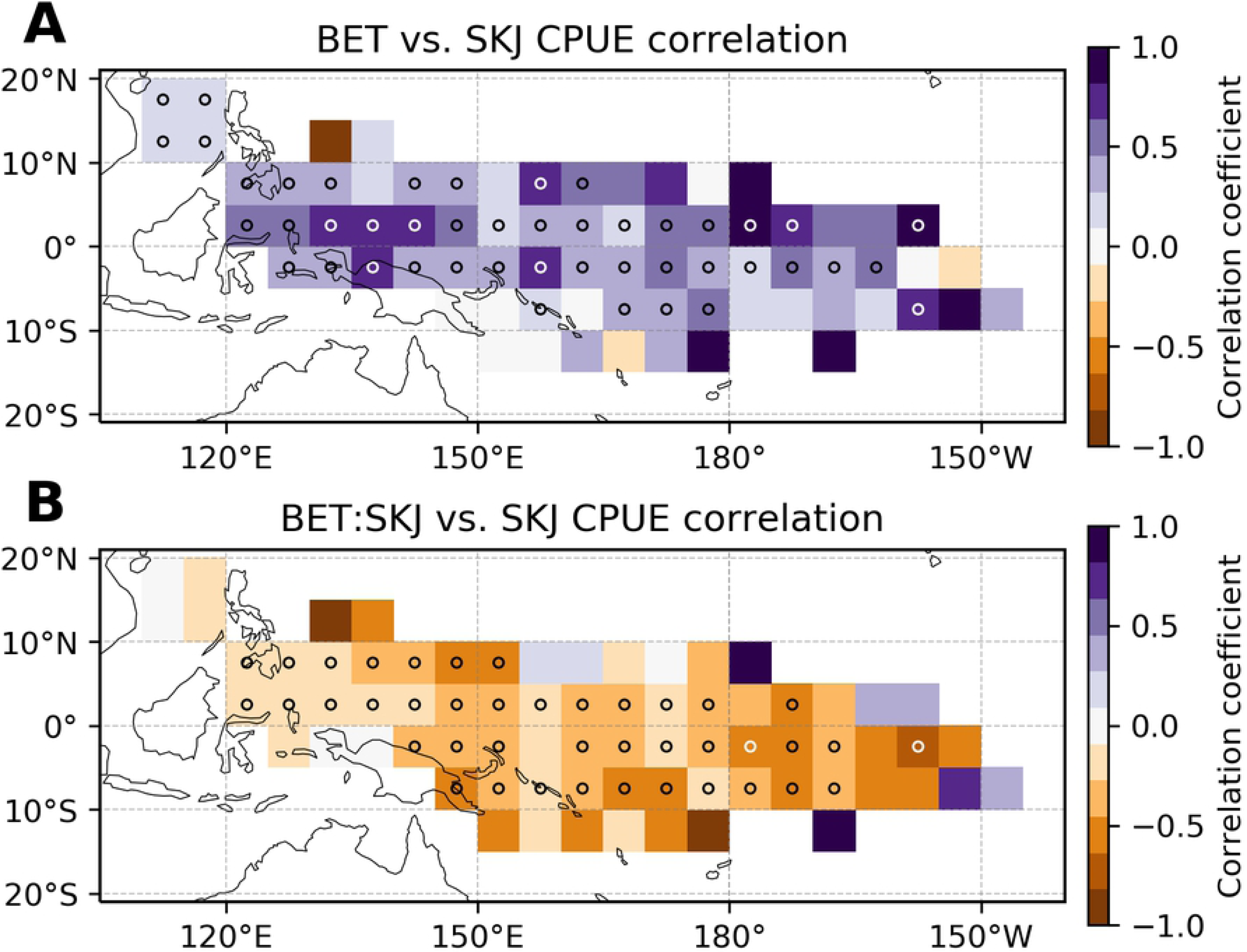
Potential lateral separability of FAD-associated skipjack and bigeye tuna. Temporal correlation coefficients between (a) monthly bigeye and skipjack catch per unit effort (CPUE), and (b) monthly bigeye-to-skipjack catch ratios and skipjack CPUE. Stippling (circles) indicates grid points where the correlation coefficient is significantly different from zero using a multiple-hypothesis-test false discovery rate of 0.1. Correlations are computed over all available data in the WCPFC purse seine catch dataset (Jan 1967 - Dec 2017). See Section 2.2.2 for further computational details.

**Table 2.**
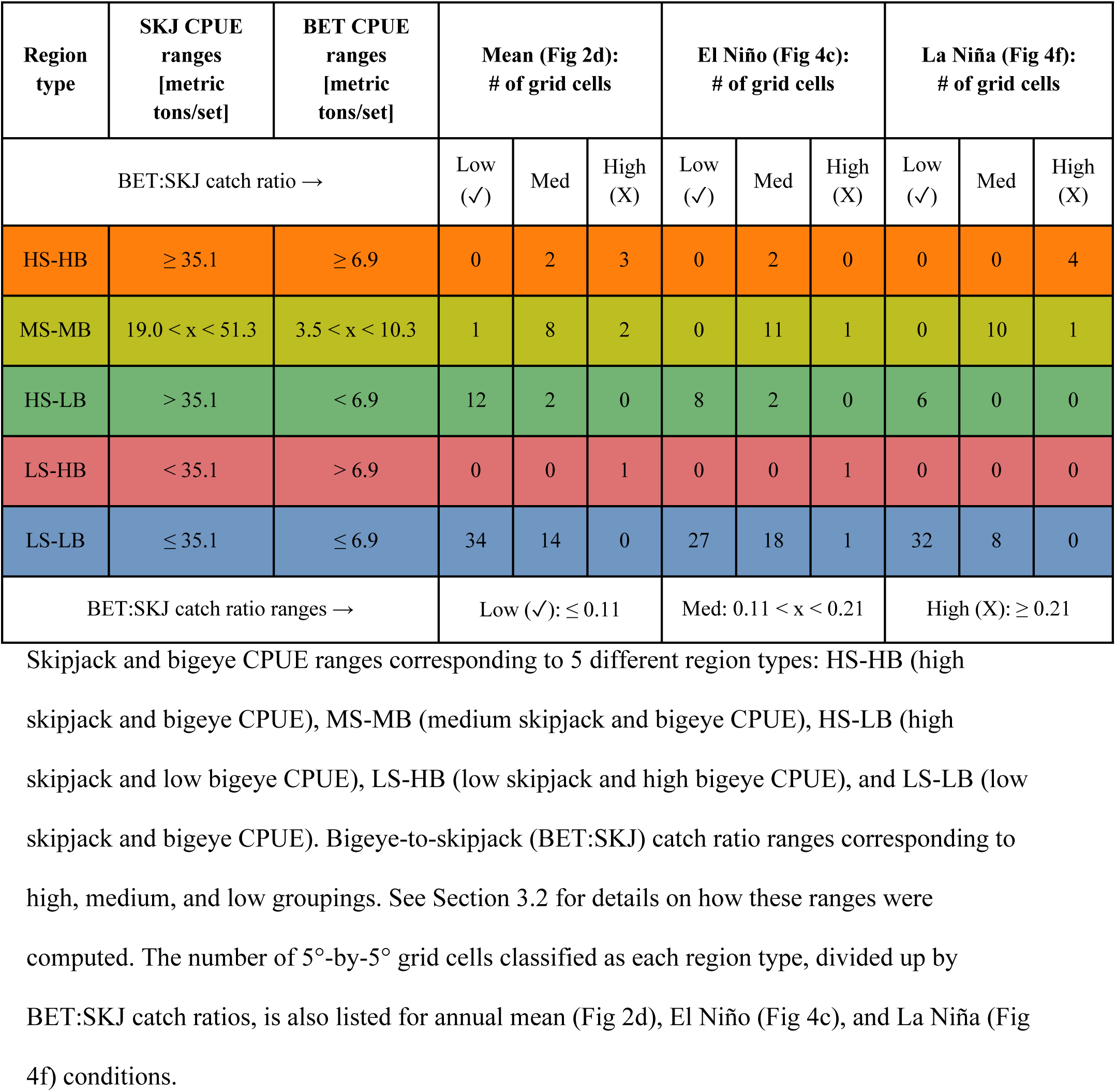
Region type classifications and occurrences.

| Region type | SKJ CPUE ranges [metric tons/set] | BET CPUE ranges [metric tons/set] | Mean (Fig 2d): # of grid cells |  |  | El Niño (Fig 4c): # of grid cells |  |  | La Niña (Fig 4f): # of grid cells |  |  |
| --- | --- | --- | --- | --- | --- | --- | --- | --- | --- | --- | --- |
| BET:SKJ catch ratio → |  |  | Low (✓) | Med | High (X) | Low (✓) | Med | High (X) | Low (✓) | Med | High (X) |
| HS-HB | $\geq 35.1$ | $\geq 6.9$ | 0 | 2 | 3 | 0 | 2 | 0 | 0 | 0 | 4 |
| MS-MB | $19.0 < x < 51.3$ | $3.5 < x < 10.3$ | 1 | 8 | 2 | 0 | 11 | 1 | 0 | 10 | 1 |
| HS-LB | $> 35.1$ | $< 6.9$ | 12 | 2 | 0 | 8 | 2 | 0 | 6 | 0 | 0 |
| LS-HB | $< 35.1$ | $> 6.9$ | 0 | 0 | 1 | 0 | 0 | 1 | 0 | 0 | 0 |
| LS-LB | $\leq 35.1$ | $\leq 6.9$ | 34 | 14 | 0 | 27 | 18 | 1 | 32 | 8 | 0 |
| BET:SKJ catch ratio ranges → | | | Low (✓): $\leq 0.11$ | | | Med: $0.11 < x < 0.21$ | | | High (X): $\geq 0.21$ | | |

#### 2.2.2 Temporal correlation maps

Temporal correlation maps (Figs 1, 8; S10-12 Figs) were created by calculating the correlation coefficient between monthly times series of two variables at each individual grid point. For all correlation coefficient maps, we also calculated the two-tailed probability, or *p*-value, that each grid point’s correlation coefficient is at least as extreme as the calculated value assuming no correlation. We deemed grid points with *p*-values below *p*_*FDR*_ (defined in Section 2.2.1 above) significantly correlated. These significantly correlated grid cells are demarcated with open circles.

#### 2.2.3 Distinguishing timescales of variability (standard deviation maps)

To separate out different sources of temporal variability in BET:SKJ catch ratios, we computed total (raw monthly) (Fig 3a), climatological (Fig 3b), anomaly-driven (Fig 3c), and ENSO anomaly-driven (Fig 3d) standard deviations in BET:SKJ catch ratios at every grid point. Total standard deviations at each grid point were computed over the entire monthly BET:SKJ time series at that grid point (Fig 3a). To compute climatological standard deviations, the climatological cycle of BET:SKJ at each grid point was first computed by averaging BET:SKJ values over each of the 12 months of the year. No lower threshold was placed on the number of years required to compute a climatological value. For example, if a given grid cell had only one valid January BET:SKJ value over the entire time series, then that data point became January’s climatological value at that grid cell. Climatological standard deviations at each grid point were then calculated over the 12-month long climatological BET:SKJ times series at that grid point (Fig 3b). These climatological standard deviations represent the amount of variability in BET:SKJ that is due to seasonality. Next, monthly anomalies of BET:SKJ were computed by subtracting the climatological cycle from the raw monthly values at each grid point. The resulting monthly anomalies were then used to compute anomaly-driven standard deviations at each grid point (Fig 3c). These monthly anomalous standard deviations represent the amount of variability in BET:SKJ due to all sources except for seasonality. Finally, ENSO anomaly-driven standard deviations were calculated over ENSO-only phases (that is, either El Niño or La Niña months) of the monthly anomalies (Fig 3d). These ENSO anomalous standard deviations represent the amount of variability in BET:SKJ due to ENSO alone (i.e., with the effects of seasonality subtracted out). The same procedures were used to create S13 Fig, which illustrates sources of temporal variability in the listed environmental variables rather than BET:SKJ catch ratios.

### 2.3 Quotient analysis

Quantification of habitat preferences was done using quotient analysis, which determines whether a given environmental condition is preferred, tolerated, or avoided by a certain species based on how frequently that species appears within that condition (e.g., [19,38,52]). The steps involved in the quotient analysis performed here are as follows:

1. Divide monthly measurements of the environmental variable under investigation into a user-specified number of equally spaced bins (30 here). Create a histogram of the given environmental variable’s values, using all measurements available at the same time and place as BET or SKJ catch data (gray bars in S8-9 Figs).
2. For each bin, compute the cumulative total SKJ or BET CPUE that corresponds with those environmental conditions (light blue bars in S8-9 Figs). For example, if sea surface temperature (SST) was the environmental variable of interest and there were a total of three 5°-by-5° monthly grid cells containing SSTs between 24.0-24.3°C over the entire CPUE-overlapped SST dataset, then we would add up all three corresponding monthly SKJ or BET CPUE values to obtain the cumulative total SKJ or BET CPUE associated with the 24.0-24.3°C SST bin. We would then repeat this procedure for all 29 other SST bins.
3. Compute the quotient, *Q*, for each bin, as follows (blue dots in S8-9 Figs): 

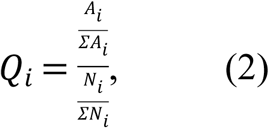

where *A*_*i*_ is total CPUE associated with bin *i* (calculated in step 2) and *N*_*i*_ is the total number of environmental measurements in bin *i* (calculated in step 1). The numerator in Eqn 2 represents the fraction of total BET or SKJ CPUE over the entire WCPFC dataset that corresponds with environmental conditions in bin *i*, while the denominator represents the fraction of all observations of the given environmental variable that fall within bin *i*. Bins with *Q*_*i*_ significantly greater than 1 contain environmental conditions that are *preferred* by a given species, while bins with *Q*_*i*_ significantly less than 1 contain conditions that are *avoided* by that species. Bins with *Q*_*i*_ that are statistically indistinguishable from 1 contain conditions that the given species merely tolerates, rather than actively prefers or avoids.
4. To determine whether *Q*_*i*_ is significantly different from 1, we compute 95% confidence intervals for the null hypothesis that the observed value of *Q*_*i*_ is drawn from a random distribution, using a resampling procedure (repeated 399 times) developed by Bernal et al. (2007) [52]. The resulting 2.5 and 97.5 percentile confidence intervals are plotted as blue dashed lines in S8-9 Figs. *Q*_*i*_ that lie outside of these confidence intervals are considered significantly different from 1.
5. Finally, to directly compare preferred skipjack and bigeye habitats, we plot their preferences (S8-9 Figs) together in one figure (Fig 7), with different colors denoting environmental conditions that are preferred, avoided, and tolerated by each species (see Fig 7 caption).

### 2.4 Code availability

The Python code and Dockerfile required to reproduce all of the figures and tables generated here can be found at https://doi.org/10.5281/zenodo.3904134.

### 2.5 Data availability

Data in the form of NetCDF files required to reproduce all of the figures and tables generated here can be found at https://doi.org/10.5281/zenodo.3904157.

## 3 Results and discussion

### 3.1 Assessment of potential bigeye-skipjack lateral separability

Monthly FAD-associated purse seine catches of skipjack and bigeye are highly positively correlated throughout the WTP (Fig 1a). Thus, in most places, BET CPUE increases whenever SKJ CPUE increases and decreases whenever SKJ CPUE decreases. This initial result suggests that the potential for lateral separation of FAD-associated skipjack and bigeye on the spatiotemporal scales investigated here may be quite limited. Correlations between monthly SKJ CPUE and BET:SKJ catch ratios tell a different story, however. As monthly SKJ CPUEs increase, fractional bigeye catches decrease throughout the WTP (Fig 1b). Thus, when skipjack are more easily captured, juvenile bigeye are caught proportionally less frequently, suggesting that the two species may be laterally separable on these spatiotemporal scales after all. (This separability is a result of the two species preferring somewhat different environmental conditions, such that they become more strongly separated when conditions are just right for skipjack, but not bigeye - see Sections 3.5 and 3.6 for details.) Having shown that lateral separation of bigeye and skipjack in the WTP is possible, we move onto quantification of where and when separation is most evident (Sections 3.2-3.5) and examine how variations in environmental conditions can alter separability (Section 3.6).

### 3.2 Mean lateral bigeye-skipjack catch separation

Side-by-side comparison of annual mean BET and SKJ CPUE maps support the notion that there is subtle but discernible lateral separation between the two species (Fig 2a-b). For instance, although both SKJ and BET CPUEs are greatest in the northeast, BET CPUE drops off dramatically east of ∼180° (Fig 2b), while SKJ CPUE remains relatively elevated within the central-eastern WTP (Fig 2a). Annual mean BET:SKJ catch ratios further demonstrate that the average degree of lateral separability between the two species is largely dependent on the exact location within the WTP (Fig 2c). Annual mean BET:SKJ catch ratios range from near zero along the southern edge of the WTP, for example, to upwards of 0.3 in the northeast corner (Fig 2c). Thus, based on catch ratios alone, the southern boundary of the WTP would be a highly desirable fishing ground for those seeking to minimize fractional bigeye bycatch, while the northeast corner may be a less desirable region.

**Fig 2.**
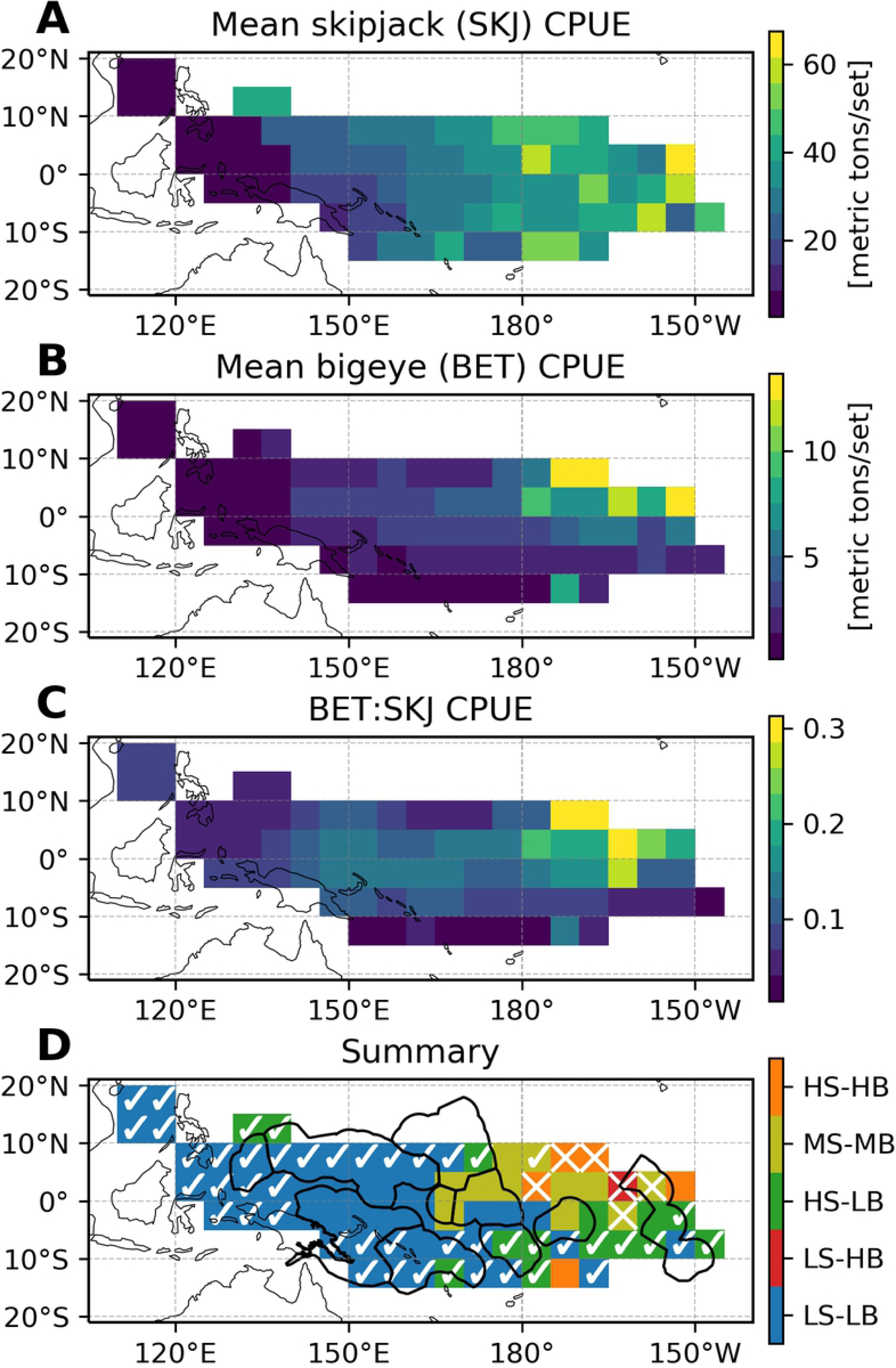
Mean FAD-associated skipjack-bigeye lateral separability. (a) Mean skipjack CPUE. (b) Mean bigeye CPUE. (c) Mean bigeye-to-skipjack catch ratio. Means are computed over all available data in the monthly WCPFC purse seine catch dataset (Jan 1967 - Dec 2017). (d) Region types based on mean skipjack (shown in a) and bigeye CPUE (shown in b) (described in detail in Section 3.2). Underlying colors correspond to region type, where S stands for skipjack, B for bigeye, H for high CPUE values, M for medium CPUE values, and L for low CPUE values. HS-LB regions, for example, exhibit high mean skipjack and low mean bigeye CPUE values. Overlain checkmarks and X’s correspond to mean bigeye-to-skipjack catch ratios (shown in c), where a checkmark denotes low (favorable) bigeye-to-skipjack catch ratios and an X denotes high (unfavorable) catch ratios. (The absence of any overlain symbol denotes moderate bigeye-to-skipjack catch ratios.) See Table 2 for CPUE and catch ratio ranges corresponding to the different region types.

However, because absolute BET and SKJ CPUE values also matter greatly in decisions of where to fish, it is most useful to examine SKJ/BET CPUE *and* BET:SKJ catch ratios at the same time to determine overall fishing desirability. We therefore combine the SKJ/BET CPUE maps in Fig 2a-b with the BET:SKJ catch ratio map in Fig 2c to produce one summary map with simultaneous information on all three factors (Fig 2d). On this summary map in Fig 2d, checkmarks denote regions with low BET:SKJ catch ratios, while X’s denote regions with high BET:SKJ catch ratios. (Regions with medium catch ratios are denoted with a lack of symbols to prevent visual clutter.) To classify catch ratios into appropriate high, medium, and low groups, we first computed minimum and maximum annual mean BET:SKJ ratios (i.e., the smallest and largest ratios in Fig 2c). We then divided the range from the minimum to maximum ratios into three equal intervals. BET:SKJ ratios within the first interval were considered low, while BET:SKJ ratios within the last (third) interval were considered high. Table 2 lists the exact cutoff values that define these intervals. To denote absolute SKJ and BET CPUE values, we also classified each grid cell on the summary map in Fig 2d into one of five region types as follows (Table 2):

1. Regions with high SKJ and BET CPUE values (high SKJ, high BET = HS-HB type; denoted in orange in Table 2 and Fig 2d)
2. Regions with medium SKJ and BET CPUE values (medium SKJ, medium BET = MS-MB type; denoted in yellow in Table 2 and Fig 2d)
3. Regions with high SKJ and low BET CPUE values (high SKJ, low BET = HS-LB type; denoted in green in Table 2 and Fig 2d)
4. Regions with low SKJ and high BET CPUE values (low SKJ, high BET = LS-HB type; denoted in red in Table 2 and Fig 2d)
5. Regions with low SKJ and BET CPUE values (low SKJ, low BET = LS-LB type; denoted in blue in Table 2 and Fig 2d)

Low, medium, and high SKJ and BET CPUE values were classified in the same way as BET:SKJ catch ratios, using four (instead of three) equally-spaced intervals. SKJ/BET CPUE values within the first two intervals were considered low, while SKJ/BET CPUE values within the last two intervals were considered high. Table 2 again lists the exact cutoff values that define these intervals. Because there can be some overlap in CPUE values between MS-MB and other region types, MS-MB regions were classified last by selecting grid cells with SKJ and BET CPUE values within their respective middle two intervals (intervals 2-3 out of 4). HS-LB regions contain the most desirable fishing grounds, where fishers can productively target skipjack while avoiding bigeye. LS-HB regions contain the least desirable fishing grounds, where fishers will catch relatively large amounts of bigeye and small amounts of skipjack. HS-HB, MS-MB, and LS-LB regions contain fishing grounds with moderate desirabilities depending on their exact BET:SKJ catch ratios. For fishers that highly value bigeye catch reduction and do not mind finding fewer skipjack, LS-LB waters may ultimately be more desirable. For fishers that must prioritize skipjack fishing efficiency and production, fishing within HS-HB waters with the lowest possible BET:SKJ catch ratios may be the right compromise.

The annual mean summary map shows the existence of all five SKJ/BET CPUE-based region types in the WTP (Fig 2d). One relatively contiguous HS-LB region occurs south of 5°S between ∼175°E-145°W (Fig 2d), where mean BET CPUE and BET:SKJ catch ratios are somewhat low (Fig 2b,c) and SKJ CPUE (Fig 2a) is relatively high. This area is thus one of the best and most desirable HS-LB regions within the WTP. In contrast, just north of this region, between ∼10°N-∼5°S and the same longitudes (∼180°-∼150°W), BET:SKJ catch ratios as well as SKJ/BET CPUE values increase substantially (Fig 2a-c), making this a less desirable MS-MB/HS-HB region (Fig 2d). LS-LB region types are primarily located west of ∼175°E, with low BET:SKJ catch ratios in the far west/north and medium BET:SKJ catch ratios in the more central WTP (Fig 2c,d). Indeed, LS-LB regions cover all grid cells west of 165°E apart from two HS-LB cells between 10°-15°N and 130°-140°E. Fishing in these LS-LB areas will likely yield small catches of skipjack as well as bigeye. The swath of ocean classified as MS-MB in the central-eastern WTP exhibits moderate SKJ and BET CPUE values, and a wide range of BET:SKJ catch ratios (Fig 2a-d). The MS-MB subregion exhibiting small to medium BET:SKJ catch ratios (165°E-165°W, 5°S-10°N) would be a reasonable area to target for those seeking to catch moderate amounts of skipjack while reducing fractional bigeye catch, compared to fishing in the high BET:SKJ catch ratio regions to the north and east (Fig 2d). Fractional bigeye catch would be higher here than in the aforementioned HS-LB regions, however. In sum, there are obvious spatial variations in the typical degree of skipjack-bigeye separation and fishing ground desirability within the WTP, with some regions exhibiting much higher (i.e., the southeast) and others exhibiting much lower (i.e., the northeast) lateral separabilities and fishing ground desirabilities.

Though these annual mean maps (Fig 2) are highly useful for understanding where lateral separation and fishing desirability is greatest on average, in reality, increased temporal resolution may be needed to make more actionable and timely decisions about where to fish. In particular, these mean maps do not provide any information on the temporal variability in the relationships between BET and SKJ CPUE caused by seasonal or ENSO-driven phenomena, both of which can greatly affect spatial distributions of tuna (e.g., [26,29]).

### 3.3 Sources of lateral separation variability

To better understand distinct sources of temporal variability in BET-SKJ separation patterns, we computed overall, climatological, anomalous, and ENSO-related standard deviations (σ) in BET:SKJ catch ratios throughout the WTP (Fig 3) (see Section 2.2.3 for more details on how these computations were done). Seasonality does not greatly affect the spatial structure of BET:SKJ catch ratios, as can be seen from the small values and distinct patterns of seasonally-driven σ’s in Fig 3b, compared to total σ’s in Fig 3a. ENSO, on the other hand, accounts for much of the temporal variability in BET:SKJ throughout the WTP, as can be seen from the large values and similar patterns of ENSO-driven σ’s in Fig 3d, compared to both monthly anomalous (Fig 3c) and total σ’s (Fig 3a). This implies that ENSO state, rather than season, is the most important determinant of where one should fish to capture skipjack but avoid bigeye. In other words, different phases of ENSO alter spatial patterns of BET:SKJ catch ratios more strongly than different seasons. In the following two sections, we analyze ENSO (Section 3.4) and seasonally-driven (Section 3.5) variations in lateral BET-SKJ separability in more detail. In Sections 3.5 and 3.6, we explain why ENSO is the more important driver of bigeye-skipjack lateral separation variability.

**Fig 3.**
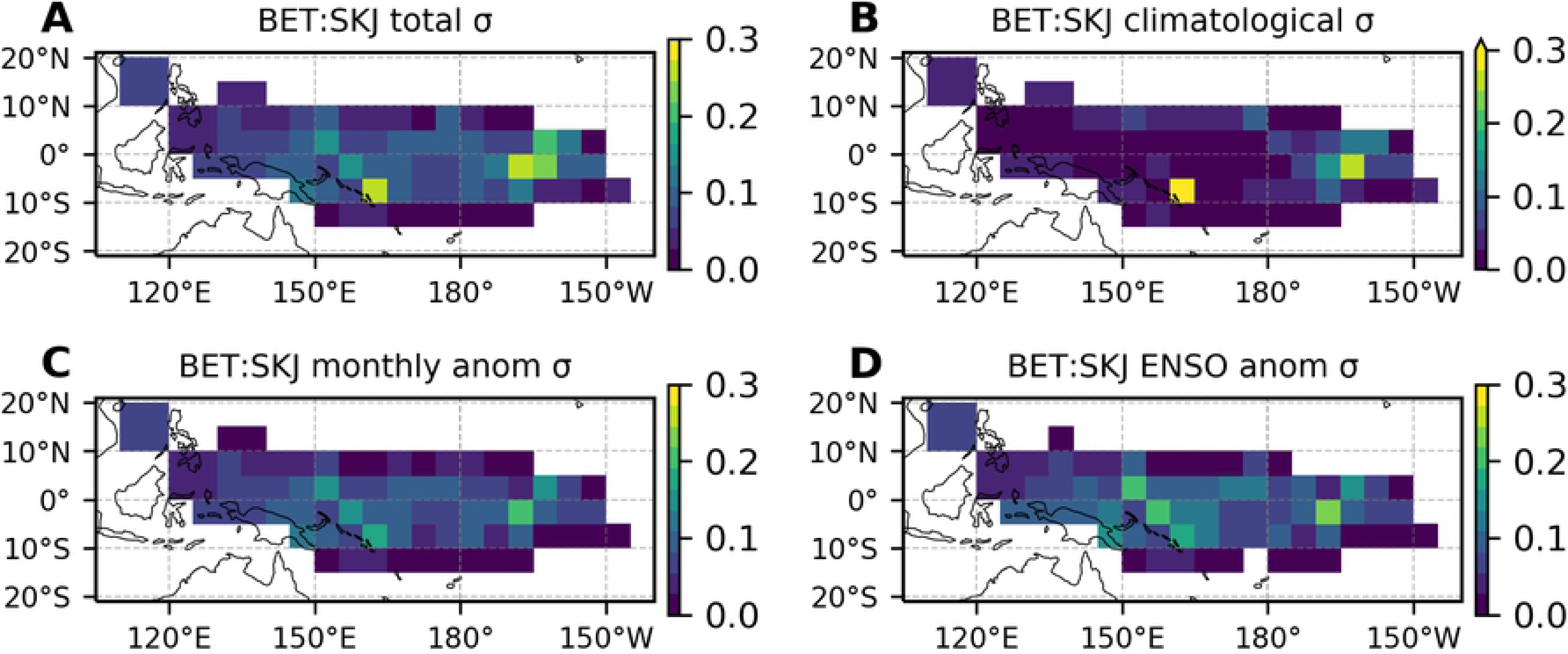
Sources of skipjack-bigeye lateral separation variability. Standard deviations (σ) of FAD-associated bigeye-to-skipjack catch ratios computed from (a) raw monthly values, (b) monthly climatologies, (c) monthly anomalies, and (d) monthly anomalies occurring during El Niño and La Niña phases. See Section 2.2.3 for further computational details.

### 3.4 ENSO-driven lateral separation variability

Spatial patterns of FAD-associated skipjack-bigeye separability vary substantially between different phases of ENSO throughout the WTP (Fig 4; S3-4 Figs) and across Party to the Nauru Agreement EEZs (PNA EEZs) (Fig 5). El Niño decreases BET:SKJ catch ratios east of ∼170°E and increases ratios west of ∼170°E, while La Niña acts in the opposite direction (Fig 4c-d; Fig 5c). The effects of El Niño and La Niña on BET:SKJ catch ratios thus oppose one another in the WTP, as is also the case for temperature, dissolved oxygen content, primary productivity, and other environmental variables (e.g., [25,53,54]).

**Fig 4.**
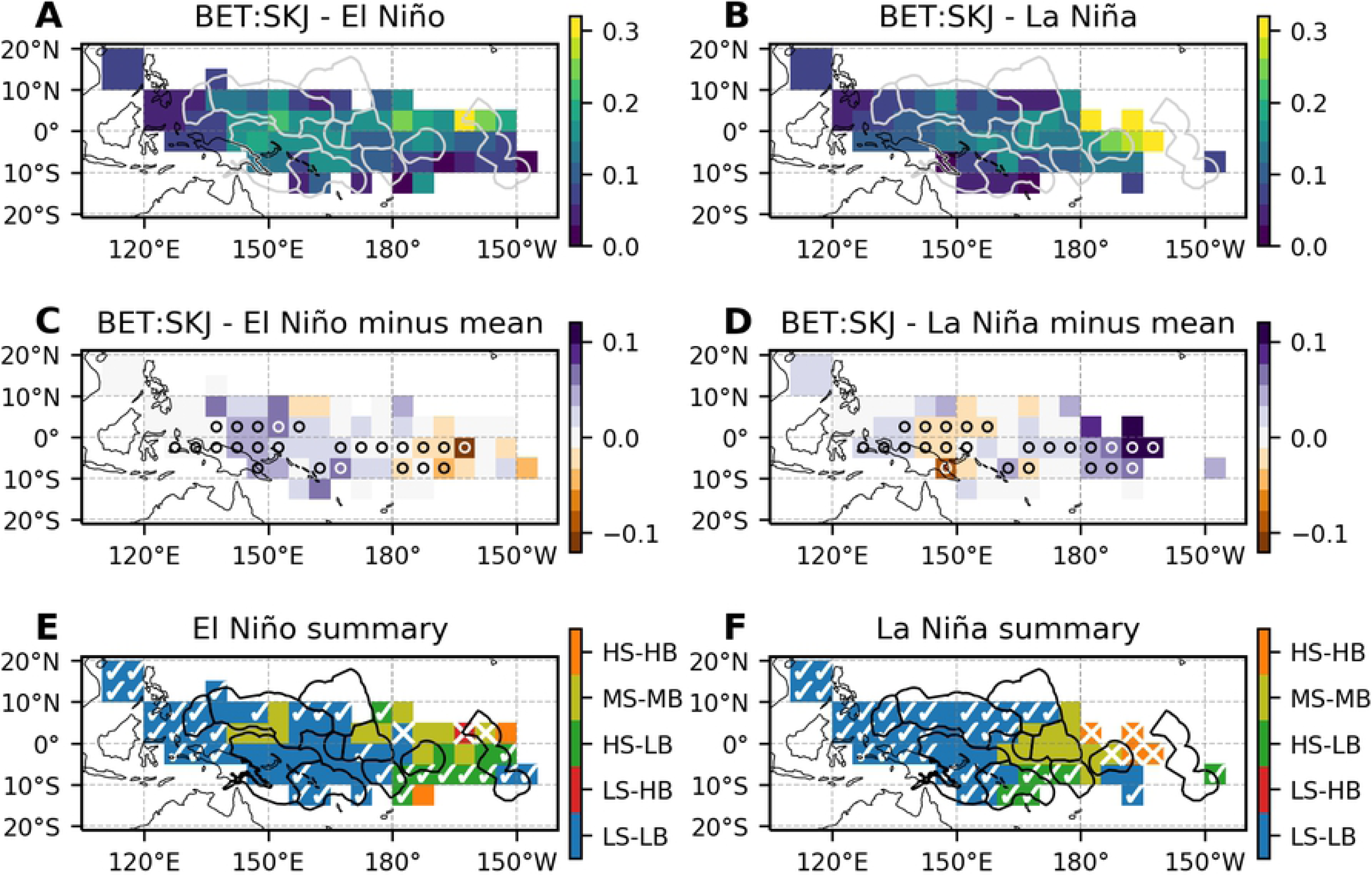
ENSO-driven skipjack-bigeye lateral separation variability. Bigeye-to-skipjack FAD-associated catch ratios averaged over (a) El Niño months and (b) La Niña months. (c, d) Same as (a, b), but with mean bigeye-to-skipjack catch ratios subtracted. Stippling (circles) indicate grid points where El Niño and La Niña composite BET:SKJ are significantly different from one another using a two-sided Wilcoxon rank-sum test and a multiple-hypothesis-test false discovery rate of 0.1 (see Section 2.2.1 for details). (e) Region types based on El Niño composite skipjack (shown in S3a Fig) and bigeye CPUE (shown in S4a Fig). Underlying colors correspond to region type, where S stands for skipjack, B for bigeye, H for high CPUE values, M for medium CPUE values, and L for low CPUE values. HS-LB regions, for example, exhibit high skipjack and low bigeye CPUE values during El Niño. Overlain checkmarks and X’s correspond to El Niño composite bigeye-to-skipjack catch ratios (shown in a), where a checkmark denotes low (favorable) bigeye-to-skipjack catch ratios and an X denotes high (unfavorable) catch ratios. (The absence of any overlain symbol denotes moderate bigeye-to-skipjack catch ratios.) See Table 2 for CPUE and catch ratio ranges corresponding to the different region types. (f) Same as (e), but based on La Niña composite maps. The light gray lines in (a,b) and black lines in (e,f) denote the exclusive economic zones (EEZs) of the eight Parties to the Nauru Agreement (PNA).

These ENSO-driven variations in skipjack-bigeye separability make the WTP region east of ∼180° longitude, where the highest annual mean BET:SKJ catch ratios occur, slightly more desirable during El Niño (i.e., BET:SKJ catch ratios are still quite high, but lower than the annual mean) and even less desirable during La Niña (i.e., BET:SKJ catch ratios are considerably higher than the annual mean) (Fig 4c-f). Kiribati’s Phoenix Island (KIR-P) EEZ is located in this region and exhibits some of the highest BET CPUE values of all PNA EEZs no matter the ENSO phase. La Niña further exacerbates the maximum bigeye CPUEs and BET:SKJ catch ratios within this EEZ (Fig 5b-c). Thus, although SKJ CPUE values are generally high here, if one’s goal is to minimize fractional BET catch, then Kiribati’s Phoenix Island EEZ should be avoided most of the time, but especially during La Niña.

**Fig 5.**
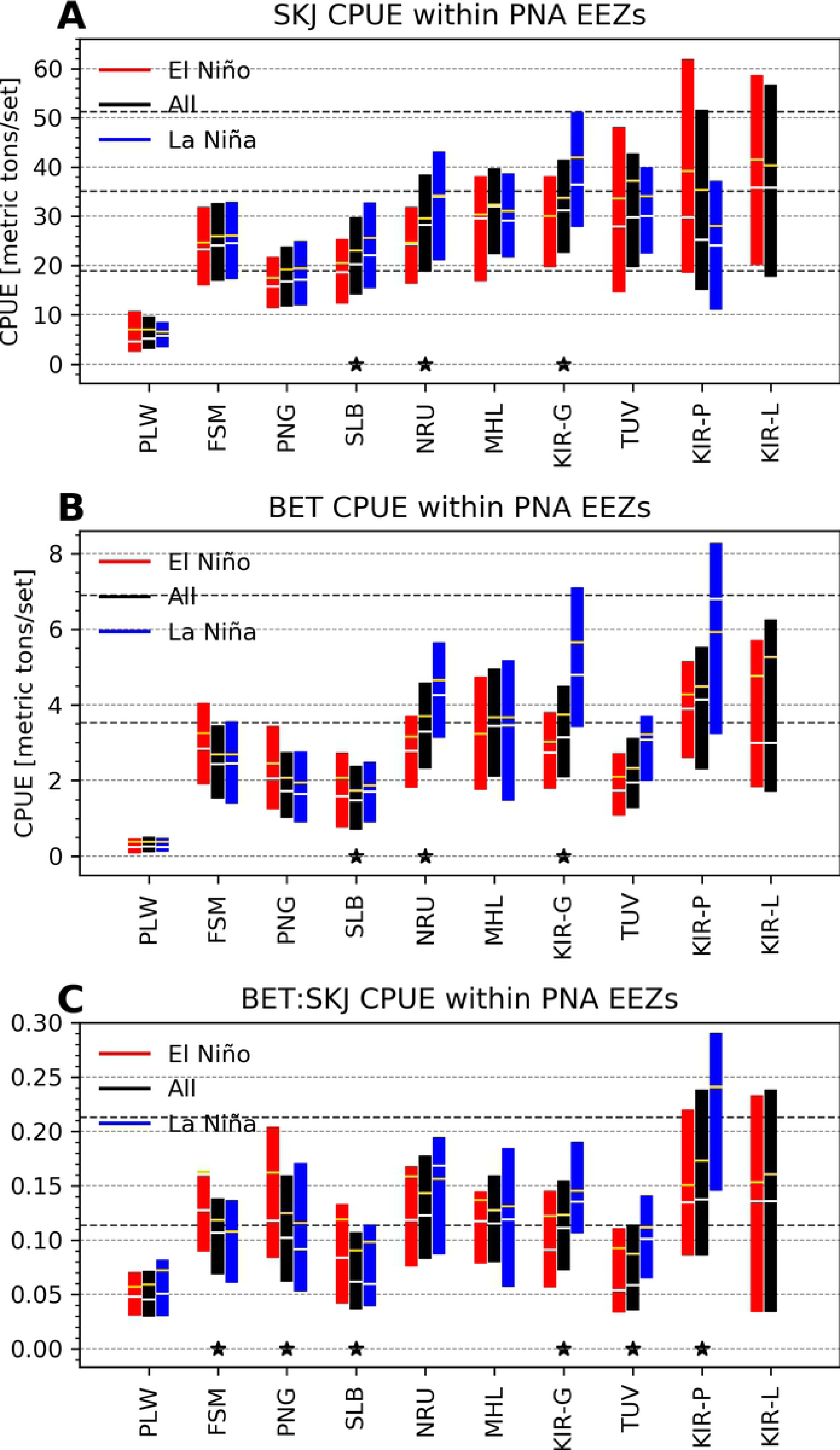
ENSO-driven lateral separation variability within Parties to the Nauru Agreement (PNA) exclusive economic zones (EEZs). Boxplot (without the whiskers; showing interquartile ranges, medians in white, and means in yellow) of FAD-associated (a) skipjack CPUE, (b) bigeye CPUE, and (c) bigeye-to-skipjack catch ratios within each PNA EEZ, composited temporally over all data (black bars), El Niño months alone (red bars), and La Niña months alone (blue bars). From left to right, the PNA countries are roughly arranged from east to west. Asterisks indicate when El Niño and La Niña composite values are significantly different from one another within a given country’s EEZ, using a two-sided Wilcoxon rank-sum test and a multiple-hypothesis-test false discovery rate of 0.1 (see Section 2.2.1 for details). Horizontal dashed lines denote the cut-off values for low, medium, and high values (see Table 2). In (c), the total number of months of BET:SKJ data available in each EEZ is listed in black below the abbreviated country names, while the number of months available during El Niño and La Niña are listed in red and blue, respectively.

Moving slightly west, between 170°E-180° and 0°-5°S, BET:SKJ catch ratios are also significantly higher during La Niña compared to El Niño (Fig 4c-d). This small 10°-longitude-by-5°-latitude region within Kiribati’s Gilbert Islands (KIR-G) EEZ thus contains desirable waters for fishers seeking to minimize fractional BET catch while simultaneously maintaining reasonably high SKJ catches during El Niño (Fig 4a,e; Fig 5a,c; S3a Figs). During La Niña, however, BET CPUEs increase substantially (Fig 5b; S4d Fig) and turn this EEZ from an LS-LB/MS-MB region with low to moderate BET:SKJ catch ratios (Fig 2d) into a fully MS-MB region with moderate BET:SKJ catch ratios (Fig 4f; Fig 5c). Fishers looking to decrease fractional BET catch may therefore try to avoid Kiribati’s Gilbert Islands EEZ during La Niña, but shift to targeting these same waters during El Niño.

Within Tuvalu’s (TUV) EEZ just south of Kiribati’s Gilbert Islands’, even though BET:SKJ catch ratios differ significantly between ENSO phases, they remain relatively low while SKJ CPUEs remain relatively high, no matter the ENSO phase (Fig 4a-b,e-f; Fig 5a,c). Tuvalu’s annual mean EEZ waters were already classified as HS-LB with low BET:SKJ catch ratios, but conditions improve even further during El Niño when BET CPUEs and BET:SKJ catch ratios are especially low (Fig 5b-c) and SKJ CPUEs do not change much (Fig 5a). Tuvalu’s EEZ waters thus regularly exhibit ideal conditions for fishers seeking productive skipjack fishing grounds containing low BET:SKJ catch rates, with conditions being especially ideal during El Niño.

Moving west across 170°E, which marks the approximate longitude where ENSO effects on BET:SKJ catch ratios switch direction, and into the region between 140-155°E and ∼5°N-10°S (largely contained within the EEZs of Micronesia and Papua New Guinea), BET:SKJ catch ratios are significantly higher during El Niño (due to increases in BET CPUE and decreases in SKJ CPUE, S3-4 Figs) and lower during La Niña (due to decreases in BET CPUE and increases in SKJ CPUE, S3-4 Figs). Papua New Guinea’s (PNG) EEZ waters exhibit especially large (though highly variable) BET:SKJ catch ratios during El Niño (Fig 5c), again driven by decreases in SKJ CPUE (Fig 5a) and increases in BET CPUE (Fig 5b). Although absolute BET CPUEs here are low all the time compared to other EEZs (including during El Niño), so are absolute SKJ CPUEs (Fig 5a-b). Papua New Guinea’s EEZ waters can thus be considered LS-LB almost all of the time, with moderate BET:SKJ catch ratios that increase during El Niño (Fig 2d; Fig 4e-f). Compared to Papua New Guinea, Micronesia’s (FSM) EEZ waters produce slightly higher (though still relatively small) absolute BET and SKJ CPUE values (Fig 5a-b), but comparable moderate BET:SKJ catch ratios (Fig 5c). BET:SKJ catch ratios also increase here during El Niño, but are slightly less variable and do not typically go as high as ratios in Papua New Guinea’s EEZ during El Niño (Fig 5c). Though relatively unproductive for skipjack, during La Niña, waters within the EEZs of Micronesia and Papua New Guinea are moderately desirable if minimization of fractional bigeye catch is the goal; this desirability is reduced by quite a bit during El Niño, however, particularly in Papua New Guinea’s EEZ, as BET:SKJ catch ratios increase significantly during this phase of ENSO.

Within Solomon Islands’ (SLB) EEZ just southeast of Papua New Guinea’s, BET:SKJ catch ratios are relatively low all the time, but are especially reduced during La Niña (Fig 5c). Absolute SKJ CPUEs tend to be higher here than in Papua New Guinea’s EEZ, but are on par with Micronesia’s EEZ to the north, particularly during La Niña (Fig 5a). Waters in and surrounding Solomon Islands’ EEZ thus exhibit some of the lowest absolute BET CPUE and BET:SKJ values of any PNA nation (Fig 5b-c), while producing low to moderate SKJ CPUEs (Fig 5a), with particularly favorable higher SKJ and lower BET CPUE values during La Niña (Fig 4f).

In sum, waters within EEZs belonging to Palau, Solomon Islands, and Tuvalu regularly exhibit the smallest BET:SKJ catch ratios of all PNA countries. SKJ CPUE is, however, quite low within Palau’s EEZ and low-to-moderate within Solomon Islands’ EEZ. In contrast, Tuvalu EEZ waters contain higher SKJ CPUE values and is therefore one of the most effective PNA EEZs to target for maximizing skipjack catch while simultaneously minimizing bigeye catch, especially during El Niño. Waters within Kiribati’s Phoenix Islands EEZ regularly exhibit the largest BET:SKJ catch ratios of the PNA countries, with especially large ratios during La Niña. Fishers looking to greatly minimize fractional bigeye catch may therefore try to avoid this region, even though SKJ CPUE is quite high here. Generally speaking, El Niño creates more favorable conditions (lower BET:SKJ catch ratios) east of ∼170°E, while La Niña creates more favorable conditions west of ∼170°E. Thus, during periods of El Niño or La Niña, the best regions to target or avoid to effectively reduce fractional bigeye catch may be quite different from those suggested by annual mean maps alone (Fig 2; Section 3.2).

### 3.5 Seasonal lateral separation variability

Seasonally-driven variations in lateral bigeye-skipjack separability are not nearly as pronounced or widespread as ENSO-driven variations (Fig 6; S5 Fig). Of the PNA EEZs, only Papua New Guinea’s exhibits significantly different BET:SKJ catch ratios between seasons (Fig 6c). Here BET:SKJ catch ratios are highest in summer and lowest in winter, though absolute differences in ratios are quite small between all seasons. In contrast, 6 of the PNA EEZs exhibited significantly different BET:SKJ catch ratios between El Niño and La Niña (Fig 5c; Section 3.4).

**Fig 6.**
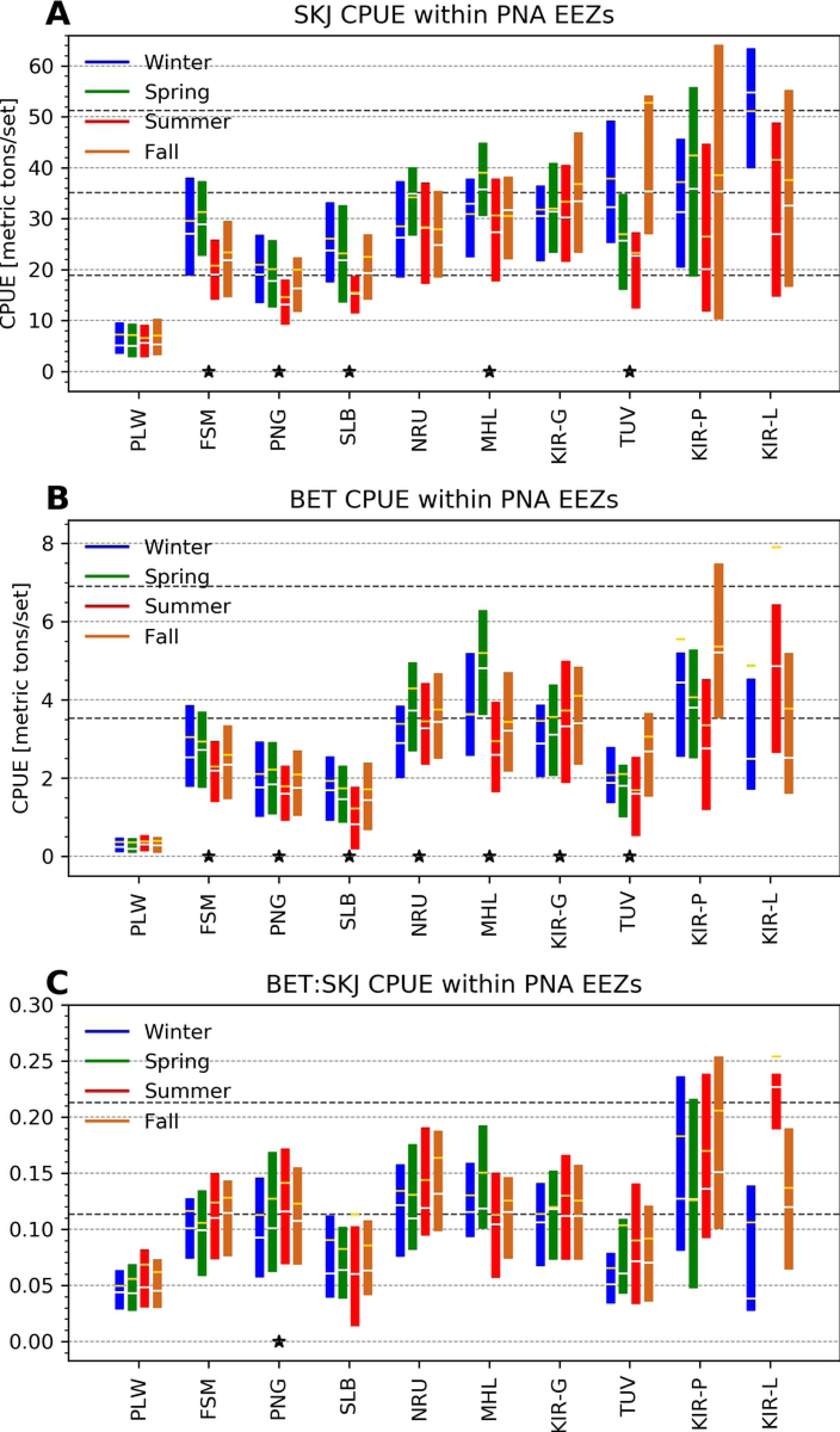
Seasonal lateral separation variability within Parties to the Nauru Agreement (PNA) exclusive economic zones (EEZs). Boxplot (without the whiskers; showing interquartile ranges, medians in white, and means in yellow) of FAD-associated (a) skipjack CPUE, (b) bigeye CPUE, and (c) bigeye-to-skipjack catch ratios within each PNA EEZ, composited temporally over the winter (December-February; blue bars), spring (March-May; green bars), summer (June-August; red bars), and autumn (September-November; brown bars) months alone. From left to right, the PNA countries are roughly arranged from east to west. Asterisks indicate when seasonal composite values are significantly different from one another within a given country’s EEZ, using a Kruskal-Wallis H-test and a multiple-hypothesis-test false discovery rate of 0.1 (see Section 2.2.1 for details). Horizontal dashed lines denote the cut-off values for low, medium, and high values (see Table 2). In (c), the number of months of BET:SKJ data available during each season in each EEZ are listed below the country names.

While significant differences in BET:SKJ catch ratios between the seasons are relatively rare and concentrated in only a couple of small regions (Fig 6c; S5 Fig), SKJ and BET CPUE values do differ significantly between the seasons in many areas throughout the WTP (Fig 6a-b; S6-7 Figs). This seemingly counterintuitive result is possible because different seasons alter spatial patterns of BET and SKJ CPUE in nearly the same way. Different phases of ENSO, on the other hand, alter spatial patterns of BET CPUE quite differently from those of SKJ CPUE (Fig 5a-b; S3-4 Figs); ENSO thus causes CPUEs for the two species to vary in different proportions or directions at any given location, ultimately leading to variable BET:SKJ catch ratios between El Niño and La Niña. The underlying physical reasons for this are discussed in Section 3.6.

In sum, the effects of ENSO on bigeye-skipjack lateral separability are more important than the effects of seasonality, though seasonality also significantly alters BET:SKJ catch ratios in some small regions within the western WTP. In these small regions, including within Papua New Guinea’s EEZ, BET:SKJ catch ratios tend to be lowest in the winter and highest in the summer, creating marginally better conditions for fractional bigeye catch minimization between December and February in this area.

### 3.6 Environmental drivers of lateral separation

Lateral separation of FAD-associated, purse seine-caught bigeye and skipjack tuna via environmental conditions alone is difficult (though not impossible) because FAD-attracted bigeye and skipjack tend to prefer and avoid very similar habitats (Fig 7; S8-9 Figs). Of the environmental variables tested here (listed in x-axes of Fig 7 subplots), only sea surface height anomalies (SSHA) can usefully distinguish between the two species’ preferred habitats over most of the WTP (Fig 8). In general, both FAD-associated skipjack and bigeye appear to prefer waters with small positive SSHA values ranging from >1 to ∼8 cm (Fig 7). However, skipjack tend to avoid negative SSHA values just below this range, while bigeye tolerate and then avoid positive SSHA values just above this range. This suggests that waters with SSHA values >∼11 cm could be effectively targeted by those seeking to minimize fractional bigeye catch. Targeting waters with positive sea surface height anomalies between ∼8 and ∼15 cm in particular would potentially minimize fractional bigeye catch while also maintaining relatively high skipjack catches. Sea surface temperatures (SST), 100-m temperatures (T_100m_), and thermocline depths (TCD) may also aid in identifying distinct skipjack and bigeye habitats, but the areas over which these environmental variables are potentially useful for this purpose are smaller than that of SSHA (S10-12 Figs). Nevertheless, SSTs around 30°C and TCDs greater than 190 m appear to be preferred by skipjack but only tolerated by bigeye, thus signifying potentially desirable temperature conditions to fish in (Fig 7; S8-9 Figs). Where T_100m_ can potentially separate the two species, skipjack prefer warmer 100-m water temperatures, while bigeye prefer cooler ones, as evidenced by the respective significant positive and negative correlation coefficients between T_100m_ and SKJ/BET CPUE (S11-12b Figs, boxed grid cells).

**Fig 7.**
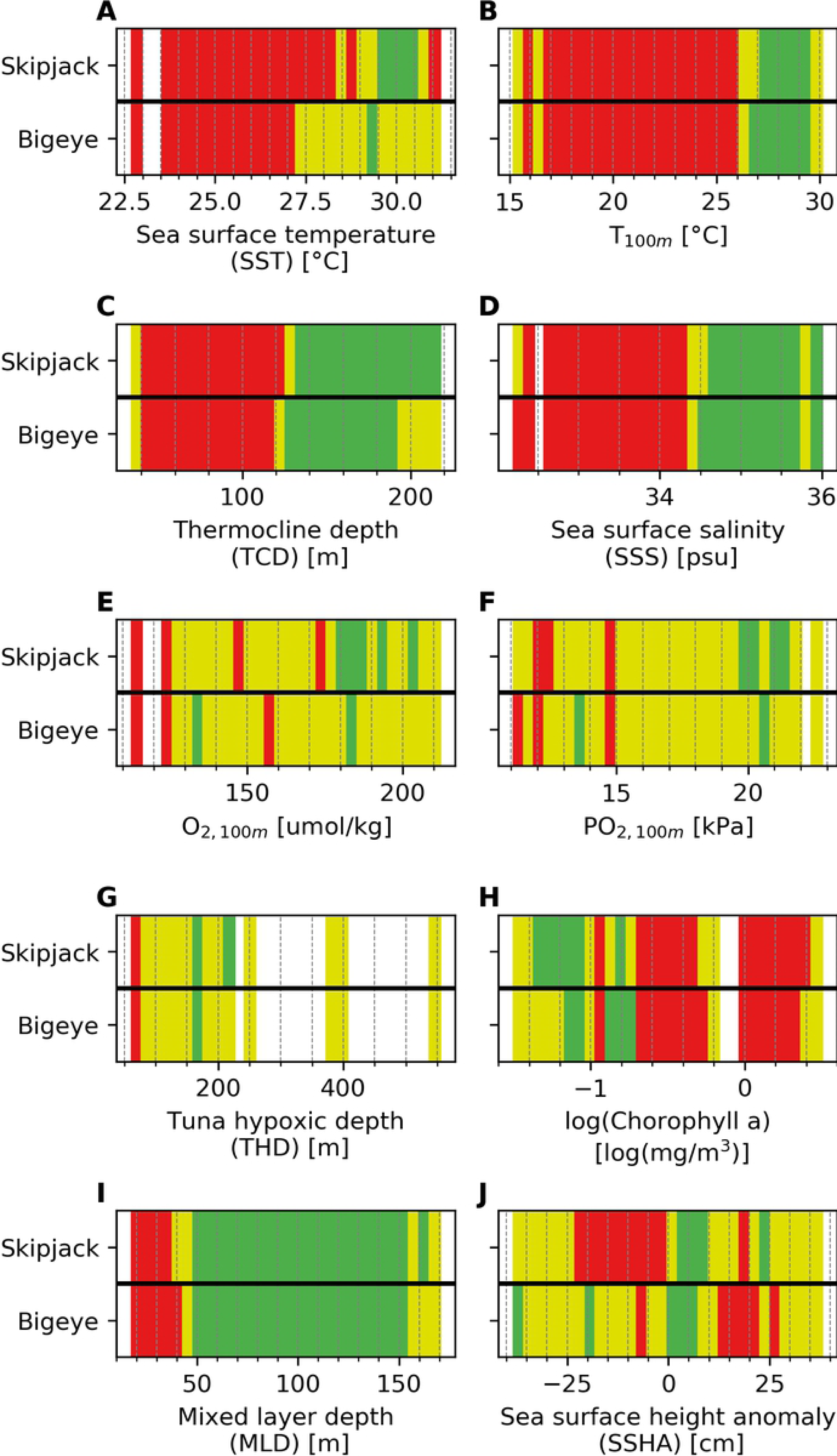
Environmental drivers of lateral FAD-associated bigeye-skipjack separability in the WTP. The color red denotes environmental conditions that the given species tends to avoid, while yellow denotes conditions that the given species tolerates and green denotes conditions that the given species prefer. The color white signifies a lack of environmental conditions in this range within the associated dataset. The top and bottom row of each subplot correspond to habitat preferences of skipjack and bigeye, respectively. See Section 2.3 for further details on how these preferences were computed.

**Fig 8.**
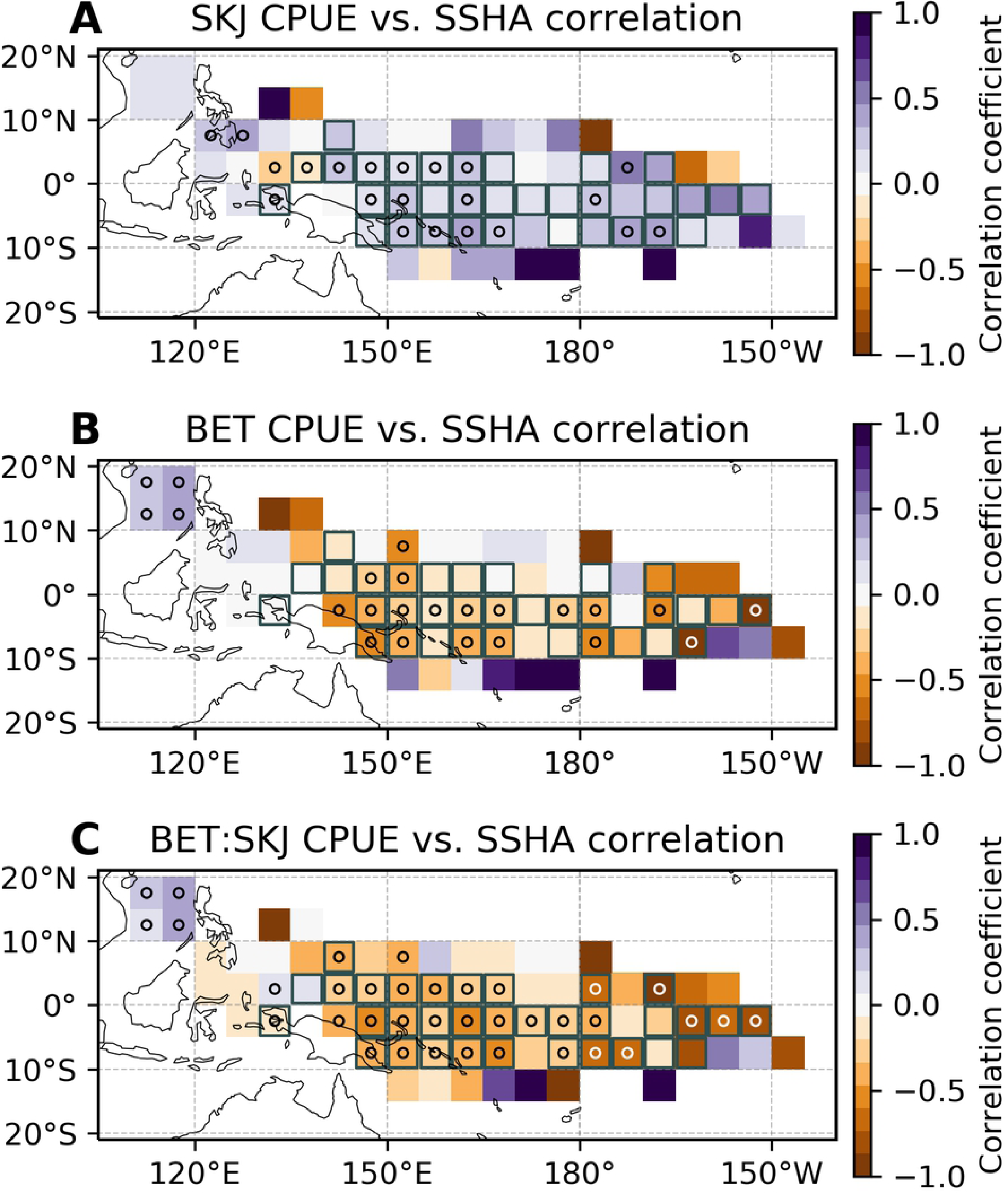
WTP regions over which sea surface height anomalies (SSHA) may be effective at laterally separating FAD-associated bigeye and skipjack. Temporal correlation coefficients between monthly SSHA and (a) skipjack CPUE, (b) bigeye CPUE, and (c) bigeye-to-skipjack catch ratios. Stippling (circles) indicates grid points where the correlation coefficient is significantly different from zero using a multiple-hypothesis-test false discovery rate of 0.1 (see Section 2.2.1 for details). Square boxes denote grid points where the following 3 criteria are met: 1.) At least one of the above maps (a-c) contains a statistically-significantly different from zero correlation coefficient; 2.) SSHA is correlated with SKJ and BET CPUEs in opposite directions (that is, the sign of the correlation coefficient is different in maps a and b above); 3.) SSHA is correlated with SKJ CPUE and BET:SKJ catch ratios in opposite directions (that is, the sign of the correlation coefficient is different in maps a and c above). Boxed grid cells thus indicate areas where a given change in SSHA conditions lead to opposite responses in skipjack and bigeye due to differential habitat preferences. Correlations are computed over all available data between Jan 1967 - Dec 2017.

These findings are in contrast to those of Hu et al. (2018) [19], who found that hypoxic layer depths, rather than temperature-related conditions and SSHAs, were most useful for laterally demarcating preferred skipjack and bigeye habitats in the ETP. There are several potential reasons for these differences. First of all, subsurface oxygen content and gradients differ greatly between the Eastern and Western Tropical Pacific [25]. In the west, oxygen is less likely to be limiting due to the presence of better oxygenated waters and much deeper hypoxic layers. Oxygen content would therefore be less likely to play an appreciable role in laterally separating bigeye and skipjack since conditions would be perfectly acceptable to both. Second of all, as in many other habitat studies, Hu et al. (2018) [19] use climatological oxygen concentrations, which may not be representative of real-time oxygen conditions, particularly during the strongest phases of El Niño or La Niña [25]. Third of all, we analyzed monthly purse seine skipjack and bigeye catches in FAD-associated sets on a 5°-by-5° horizontal grid, while Hu et al. (2018) [19] analyzed monthly purse seine catches in all set types on a 1°-by-1° horizontal grid. Because of these differences in set type and/or spatial resolution, results from our two studies may not be directly comparable. Lastly, because observations of oxygen in the WTP are relatively sparse compared to those of temperature and sea surface height anomalies, it is possible that there was simply not enough data to differentiate between the oxygen conditions preferred by skipjack and bigeye. With more complete observations, we may find that subsurface oxygen conditions can also serve to effectively separate FAD-associated bigeye and skipjack in the WTP.

In Sections 3.3 and 3.5, we noted that different ENSO phases alter spatial patterns in BET:SKJ catch ratios more strongly than different seasons in the WTP. This is because ENSO-driven changes in environmental conditions that separate FAD-associated bigeye and skipjack (SSHA, T_100m_, TCD, and SST) are more extreme than those driven by changing seasons here (S13 Fig). These larger ENSO-driven habitat variations are more likely to create conditions that reach the limit of one species’ preferences or avoidances, while entering another species’ preferred or avoided range. In contrast, the much smaller seasonal changes in SSHA, T_100m_, TCD, and SST tend to drive bigeye and skipjack abundances in the same direction, leading to variations in both species’ CPUEs, but not in resultant BET:SKJ catch ratios.

In sum, skipjack and bigeye are not easily laterally separated by differences in habitat preferences alone, but SSHA conditions may be most useful in demarcating their preferred environments. SST, T_100m_, and TCD may also play secondary roles in distinguishing habitats preferred/avoided by FAD-associated skipjack and bigeye within the WTP. Temperature-related environmental conditions thus appear to be more important drivers of lateral skipjack-bigeye separability in the WTP, while oxygen appears to be more important in the ETP [19]. Larger ENSO-driven changes in environmental conditions are able to better separate skipjack and bigeye compared to seasonally-driven changes, given the two species’ somewhat distinct habitat preferences.

## 4 Conclusions

Incidental capture of juvenile bigeye tuna in FAD-associated purse seine fisheries targeting skipjack has contributed significantly to the degradation of bigeye stocks in the Western Tropical Pacific (WTP). However, efforts to reduce bigeye catch should also prioritize maintenance of skipjack catch at current levels. Here we examined spatial patterns in FAD-associated purse seine bigeye and skipjack CPUE, as well as bigeye-to-skipjack catch ratios, and quantified how these spatial patterns vary between different seasons and phases of ENSO. We also examined the environmental conditions that are most efficient at laterally separating bigeye and skipjack. We find that FAD-associated bigeye and skipjack CPUE covary tightly throughout the WTP on the spatiotemporal scales examined here (5°-by-5°, monthly). There are, however, significant variations in this separability both spatially and temporally. Temporally, El Niño phases create more desirable conditions (lower bigeye-to-skipjack catch ratios) east of ∼170°E longitude, while La Niña phases create more desirable conditions west of ∼170°E. The effects of these ENSO cycles on bigeye-skipjack separability were found to be more important than the effects of seasonality throughout the WTP. Spatially, waters within the EEZs belonging to Palau, Solomon Islands, and Tuvalu regularly exhibit some of the smallest bigeye-to-skipjack catch ratios in the WTP. Tuvalu’s EEZ waters also exhibit relatively high skipjack CPUEs and is therefore one of the most desirable EEZs to target when seeking to simultaneously minimize bigeye while maximizing skipjack catch, particularly during El Niño. In contrast, waters within Kiribati’s Phoenix Islands EEZ regularly exhibit some of the largest bigeye-to-skipjack catch ratios, with especially large ratios during La Niña. Fishers looking to minimize fractional bigeye catch may therefore want to avoid this region, even though skipjack CPUE is quite high here. Skipjack and bigeye are not easily laterally separated by differences in habitat preferences alone, but real-time maps of sea surface height anomalies (available from https://marine.copernicus.eu/ and others) may be useful in demarcating their different environments. Sea surface temperatures, temperatures at 100 m, and thermocline depths may also be helpful in distinguishing between habitats preferred by skipjack and bigeye.

Two caveats of this study to keep in mind are that the FAD-associated purse seine catches relied upon here may sometimes be misreported or misidentified [55], and CPUE values are not always a representative proxy for abundance [56]. Nevertheless, we show that there are significant spatiotemporal variations in FAD-associated bigeye-skipjack purse seine catch separability that should be taken into account when determining where to fish or to implement FAD set restrictions.

## Supporting information

**S1 Fig. Number of purse seine sets of different types in the WCPFC catch and effort dataset.** Total number of (a) natural log/debris associated sets, (b) drifting fish-aggregating device (FAD) associated sets, (c) anchored FAD associated sets, (d) all associated sets (natural log/debris, drifting FAD, and anchored FAD combined), (e) unassociated sets, and (f) other types of sets in the WCPFC purse seine catch and effort dataset.

**S2 Fig. Number of monthly data points available for skipjack/bigeye CPUEs and corresponding environmental conditions.** (a, b) Total number of months of data on skipjack and bigeye CPUE available in the WCPFC purse seine catch dataset. (c-l) Total number of months of available environmental data overlapping with available months of both skipjack and bigeye CPUE. See Table 1 for further information on each environmental variable’s dataset.

**S3 Fig. ENSO-driven skipjack CPUE variability.** Skipjack CPUE averaged over (a) El Niño months and (b) La Niña months. (c, d) Same as (a, b), but with mean SKJ CPUE subtracted. Stippling (circles) indicate grid points where El Niño and La Niña composite SKJ CPUE are significantly different from one another using a two-sided Wilcoxon rank-sum test and a multiple-hypothesis-test false discovery rate of 0.1 (see Section 2.2.1 for details). The light gray lines in (a,b) denote the exclusive economic zones (EEZs) of the eight Parties to the Nauru Agreement (PNA).

**S4 Fig. ENSO-driven bigeye CPUE variability.** Bigeye CPUE averaged over (a) El Niño months and (b) La Niña months. (c, d) Same as (a, b), but with mean BET CPUE subtracted. Stippling (circles) indicate grid points where El Niño and La Niña composite BET CPUE are significantly different from one another using a two-sided Wilcoxon rank-sum test and a multiple-hypothesis-test false discovery rate of 0.1 (see Section 2.2.1 for details). The light gray lines in (a,b) denote the exclusive economic zones (EEZs) of the eight Parties to the Nauru Agreement (PNA).

**S5 Fig. Seasonally-driven skipjack and bigeye lateral separation variability.** (Column 1) Bigeye-to-skipjack catch ratios averaged over winter, spring, summer, and autumn months. (Column 2) Same as (Column 1), but with mean bigeye-to-skipjack catch ratios subtracted. Stippling (circles) in (Column 2) indicate grid points where seasonal composite BET:SKJ values are significantly different from one another using a Kruskal-Wallis H-test and a multiple-hypothesis-test false discovery rate of 0.1 (see Section 2.2.1 for details). (Column 3) Region types based on seasonal mean skipjack (shown in S3a Fig) and bigeye CPUE (shown in S4a Fig). Underlying colors correspond to region type, where S stands for skipjack, B for bigeye, H for high CPUE values, M for medium CPUE values, and L for low CPUE values. HS-LB regions, for example, exhibit high seasonal mean skipjack and low seasonal mean bigeye CPUE values. Overlain checkmarks and X’s correspond to seasonal mean bigeye-to-skipjack catch ratios (shown in Column 1), where a checkmark denotes low (favorable) bigeye-to-skipjack catch ratios and an X denotes high (unfavorable) catch ratios. (The absence of any overlain symbol denotes moderate bigeye-to-skipjack catch ratios.) See Table 2 for CPUE and catch ratio ranges corresponding to the different region types. The light gray lines in Column 1 and black lines in Column 3 denote the exclusive economic zones (EEZs) of the eight Parties to the Nauru Agreement (PNA).

**S6 Fig. Seasonally-driven skipjack CPUE variability.** (Column 1) Skipjack CPUE averaged over winter, spring, summer, and autumn months. (Column 2) Same as (Column 1), but with mean skipjack CPUE subtracted. Stippling (circles) in (Column 2) indicate grid points where seasonal composite SKJ CPUE values are significantly different from one another using a Kruskal-Wallis H-test and a multiple-hypothesis-test false discovery rate of 0.1 (see Section 2.2.1 for details). The light gray lines in Column 1 denote the exclusive economic zones (EEZs) of the eight Parties to the Nauru Agreement (PNA).

**S7 Fig. Seasonally-driven bigeye CPUE variability.** (Column 1) Bigeye CPUE averaged over winter, spring, summer, and autumn months. (Column 2) Same as (Column 1), but with mean bigeye CPUE subtracted. Stippling (circles) in (Column 2) indicate grid points where seasonal composite BET CPUE values are significantly different from one another using a Kruskal-Wallis H-test and a multiple-hypothesis-test false discovery rate of 0.1 (see Section 2.2.1 for details). The light gray lines in Column 1 denote the exclusive economic zones (EEZs) of the eight Parties to the Nauru Agreement (PNA).

**S8 Fig. Quotient analysis for associated purse seine-caught skipjack habitat preferences.** See Section 2.3 for details.

**S9 Fig. Quotient analysis for associated purse seine-caught bigeye habitat preferences.** See Section 2.3 for details.

**S10 Fig. WTP regions over which different environmental variables may be effective at laterally separating bigeye and skipjack.** Temporal correlation coefficients between monthly BET:SKJ catch ratios and the environmental variables shown. Stippling (circles) indicates grid points where the correlation coefficient is significantly different from zero using a multiple-hypothesis-test false discovery rate of 0.1 (see Section 2.2.1 for details). Square boxes denote grid points where the following 3 criteria are met: 1.) At least one of the corresponding maps in S10-12 Figs contains a statistically-significantly different from zero correlation coefficient; 2.) The given environmental variable is correlated with SKJ and BET CPUEs in opposite directions (that is, the sign of the correlation coefficient is different on corresponding maps in S11-12 Figs); 3.) The given environmental variable is correlated with SKJ CPUE and BET:SKJ catch ratios in opposite directions (that is, the sign of the correlation coefficient is different on corresponding maps in S10-11 Figs). Boxed grid cells thus indicate areas where a given change in the environmental variable at hand leads to opposite responses in skipjack and bigeye due to differential habitat preferences. Correlations are computed over all available data between Jan 1967 - Dec 2017.

**S11 Fig. Correlations between environmental conditions and skipjack CPUE.** Temporal correlation coefficients between monthly SKJ CPUE values and the environmental variables shown. S10 Fig’s legend applies here as well.

**S12 Fig. Correlations between environmental conditions and bigeye CPUE.** Temporal correlation coefficients between monthly BET CPUE values and the environmental variables shown. S10 Fig’s legend applies here as well.

**S13 Fig. Sources of environmental variability.** Standard deviations of sea surface height anomalies (SSHA) computed from (a) monthly anomalies occurring during El Niño/La Niña phases and (b) from monthly climatologies. (c) Difference between standard deviations computed from ENSO-driven monthly anomalies and monthly climatologies (that is, a minus b). See Section 2.2.3 for further computational details. (d-f) Same as (a-c), but for 100-m temperatures (T_100m_). (g-i) Same as (a-c), but for thermocline depths (TCD). (j-l) Same as (a-c), but for sea surface temperatures (SST).

